# Patterns of cancer somatic mutations predict genes involved in phenotypic abnormalities and genetic diseases

**DOI:** 10.1101/120121

**Authors:** Paolo Provero, Ivan Molineris, Dejan Lazarevic, Davide Cittaro

## Abstract

Genomic sequence mutations in both the germline and somatic cells can be pathogenic. Several authors have observed that often the same genes are involved in cancer when mutated in somatic cells and in genetic diseases when mutated in the germline. Recent advances in high-throughput sequencing techniques have provided us with large databases of both types of mutations, allowing us to investigate this issue in a systematic way. Here we show that high-throughput data about the frequency of somatic mutations in the most common cancers can be used to predict the genes involved in abnormal phenotypes and diseases. The predictive power of somatic mutation patterns is largely independent of that of methods based on germline mutation frequency, so that they can be fruitfully integrated into algorithms for the prioritization of causal variants. Our results confirm the deep relationship between pathogenic mutations in somatic and germline cells, provide new insight into the common origin of cancer and genetic diseases and can be used to improve the identification of new disease genes.

## 1 Introduction

Cancer has been called a disease of the genome since in most cases it is initiated by mutations occurring in somatic cells leading to uncontrolled proliferation and eventually to metastatic invasion of other tissues. On the other hand many diseases, both rare and common, can be caused or favored by mutations in the germline genome. It is thus natural to ask to what extent the same mutations can be associated to cancer and genetic diseases when occurring respectively in somatic or germline cells.

Indeed many cases are known of genes involved in both types of diseases: for example Rasopathies [1] are a family of developmental diseases caused by germline mutations in genes of the Ras/MAPK pathway, which is also recurrently mutated in many cancer types [2]. A recent review [3] pointed out the role of mutations of chromatin remodelers in both cancer and neurodevelopmental disorders. However, to our knowledge, the extent to which the mutational spectrum of cancer and genetic disorders overlap has never been investigated in a systematic way.

Recent development in sequencing technologies allow the determination of mutations in patients in a fast and cost-effective way, especially when the sequencing is limited to exons, so that sequencing is now routinely used as a diagnostic and prognostic tool in both genetic diseases and cancer. These development have also allowed the creation of large databases of mutations including many thousands of individuals, providing us with the means to investigate the relationship between somatic and germline pathogenic mutations in a systematic and statistically controlled way.

We thus decided to investigate whether patterns of somatic mutations detected in cancer samples contain information that can be used to predict the involvment of genes in genetic diseases. We chose to tackle the issue in a machine-learning framework, that is to reframe the question as whether it is possible to predict the involvement of a gene in a genetic disease (or more generally an abnormal phenotype due to germline mutations) using the frequency of its somatic mutations in a set of common cancers. In this way we can take advantage of the statistical methods developed in machine learning to accurately quantify the predictive power of the model, and to determine whether such cancer-based predictors can provide new information when combined with more traditional disease-gene prioritization methods, based for example on the frequency spectrum of germline mutations.

## 2 Results

### 2.1 Frequency of somatic mutations in cancer predicts involvement in abnormal phenotypes

We obtained from the TCGA [4] project the frequency of somatic, exonic, non-silent mutations for 18499 protein-coding genes in 29 cancers. From the Human Phenotype Ontology (HPO) [5] we obtained the association between 1007 phenotypes and 3229 genes (we considered only phenotypes with more than 50 associated genes to avoid problems in fitting logistic models. Moreover we limited the analysis to HPO terms classified as “abnormal phenotypes” but not as “neoplasms”).

We first considered, for each gene, the total number of somatic mutations observed in cancer, summed over all 29 cancer types (Total Somatic Mutations - TSM in the following). For each of the 1007 phenotypes we fitted a univariate logistic model in which the regressor is the TSM and the regressed binary variable is the involvment of the gene in the phenotype according to the HPO annotation. The performance of the models were evaluated by their Area Under the Receiver Operating Characteristic (ROC) curve (AUC), and their statistical significance by a Mann-Whitney *U test*.

We found that for most abnormal phenotypes (658 out of 1007) the TSM is indeed a significant predictor of gene involvment (Bonferroni-corrected *P* < 0.05). For almost all of these (654 out of 658) the coefficient of the logistic model is positive, that is TSM is *positively* associated to involvement in the disease. The exceptions are phenotypes related to exercise intolerance and mitochondrial function (*Exercise intolerance*, *Abnormality of the mitochondrion*, *Abnormality of mitochondrial metabolism* and *Lactic acidosis*): the genes involved in these phenotypes are characterized by low TSM, possibly due to negative selection operating, in tumors, on these genes. These results do not depend strongly on the precise choice of the regressor variable, as essentially identical numbers are obtained using the total frequency of somatic mutations (in which the number of mutations for each cancer type is divided by the number of samples available in the TCGA), or by log-transforming total occurrences or frequencies before summing over cancer types.

### 2.2 Comparison with germline-based predictors

Several predictors of disease genes have been developed, often based, as it is natural, on germline mutation frequencies. For example the “Residual Variation Intolerance Score” (RVIS) derived in [6] uses data from a large sample of whole exome sequencing (of germline DNA) data to determine a score predicting the likelihood of a gene to be involved in a disease. Similarly the authors of [7] estimated for all genes the probabilities of being tolerant of both homozygous or heterozygous loss-of-function variants. We thus compared the power of TSM in predicting HPO annotations to these germline-based scores: it turns out that TMS is a slightly *more* powerful predictor than either germline-derived indicators, as shown in Fig. 1.

**Figure 1:**
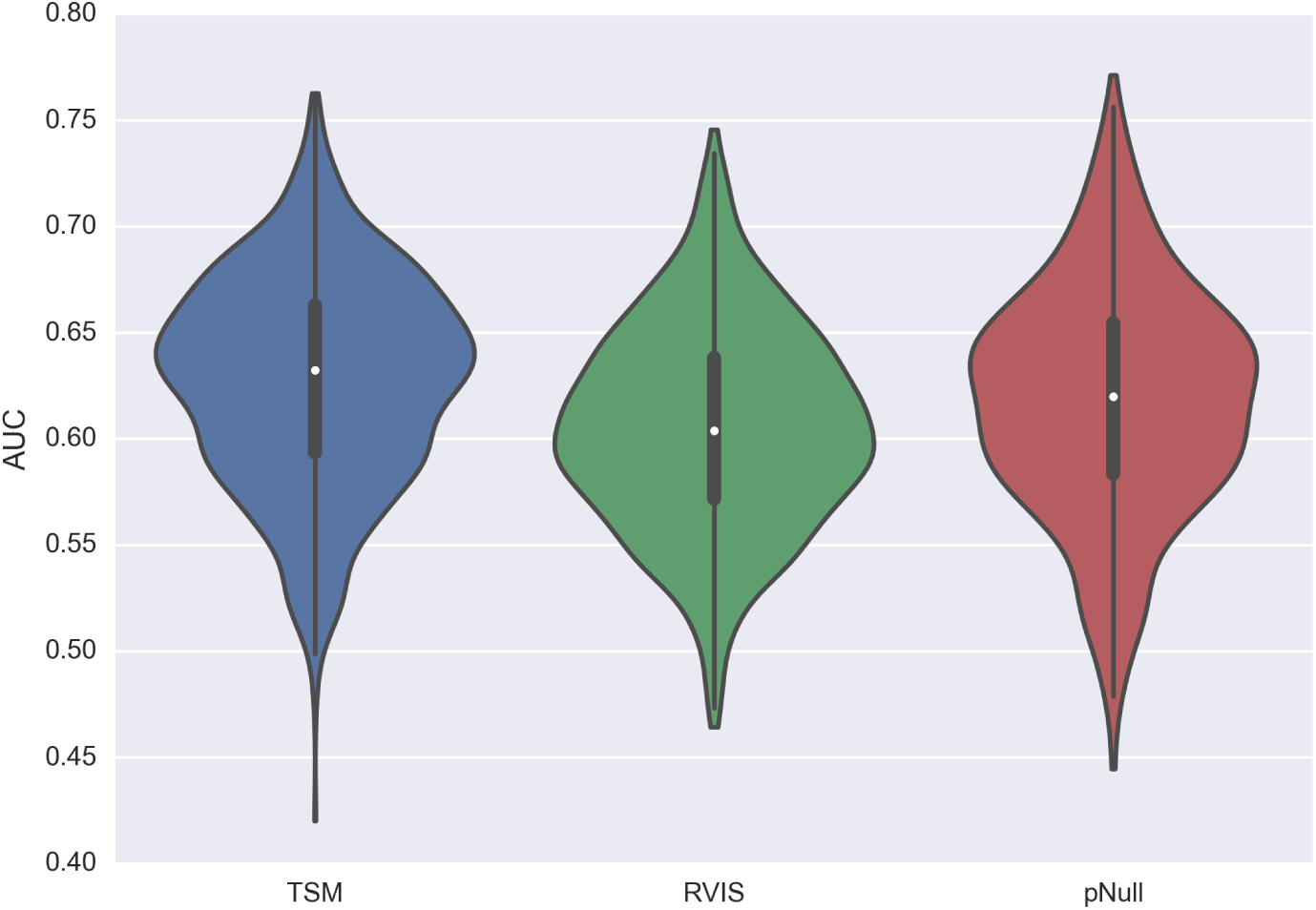
Distribution of AUCs obtained for 1007 phenotypes using TSM, RVIS [6] and the probability of being tolerant of loss-of-function variants [7]

### 2.3 TSM and germline-based indicators are independent disease gene predictors

The fact that both TSM and germline-based indicators are useful disease gene predictors suggest that their integration could attain even higher predictive power, as long as the information they provide is at least partially independent. To verify this we fitted bivariate logistic models using TSM and each germline indicator as regressors. The bivariate models achieved, as expected, higher AUC values than the univariate ones. For example the bivariate model using TSM and pNull as regressors was a significant disease gene predictor for 724 HPO phenotypes. More importantly, for 675 of these phenotypes the coefficient of at least one of the regressors was significantly different from zero (*P* < 0.05), implying that the two regressor provide independent information about the involvement in disease. These results imply that somatic mutation frequency profiles may be usefully integrated with germline-based indicators in predicting disease genes.

### 2.4 The predictive power of TSM in specific cancer types

Having concluded that TSM is a useful predictor of disease genes, we specifically asked which cancer types were more predicitve of involvment in which specific diseases. To this aim we fitted univariate logistic models as above, but using as regressors the TSM for each cancer type separately. The results are shown in Fig. 2, which represents the AUCs achieved by each cancer-type-specific TSM for the top-level disease classes, i.e. the terms that are direct descendants of “Phenotypic abnormality” in the HPO tree.

**Figure 2:**
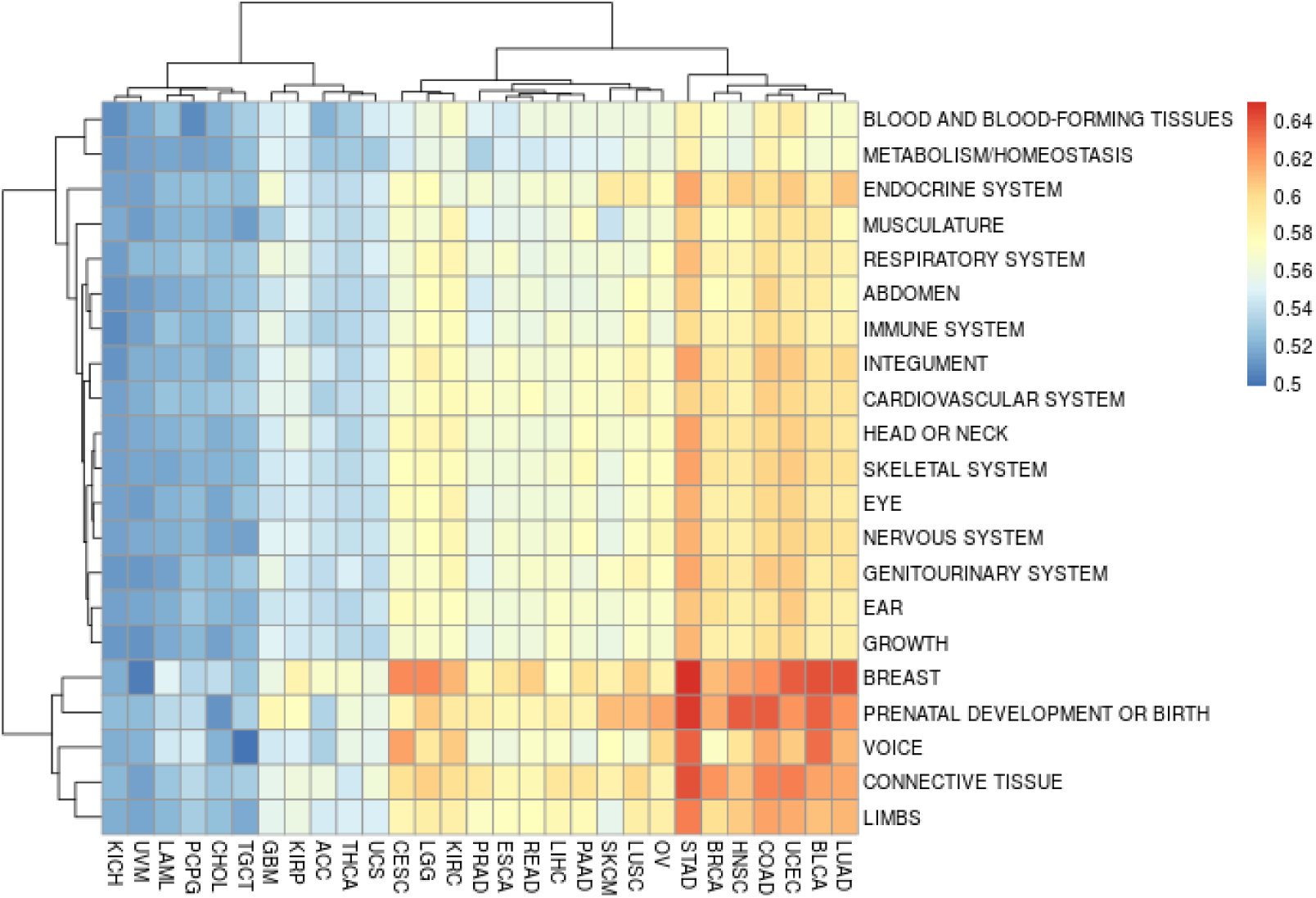
Heatmap representing the AUC achieved by TSM models based on individual tumor types as predictor of genes involved in the most general HPO phenotypes

We note a cluster of 7 cancer types with strong predictive power across most disease classes. At the opposite spectrum, a cluster of 11 cancer types show almost no predictive power on any disease class. Finally a cluster of 11 cancer types shows predictive power only for a subset of disease classes.

Somehow surprisingly, the high-predictive cancer types include stomach, lung and colon adenocarcinoma, that according to [8] are among the cancer types where the spectrum of variant frequency best agrees with a model of *neutral* evolution, thus casting doubt on the *functional* significance of mutations found in these tumors. Symmetrically, the least predictive cancer types include glioblastoma and thyroid carcinoma, among the ones in which the evolution is farthest from neutral according to [8]. It must be noted however that the TSM we use was derived by the TCGA consortium using filters on frequency such that most of the mutations we consider are clonal or in any case above the frequency range for which the *1/f* distribution characteristic of neutral evolution is observed in [8]. In addition, the number of somatic mutations is higher for cancer types showing neutral evolution (see Supplementary Figure 2); in this sense, cancer under neutral evolution model may explore the space of possible mutations in a broader way, compared to other tumor types and hence be better predictors.

Table 1 shows top 20 abnormal phenotypes in term of AUC, together with the tumor type that provides the best predictor. The same data for all abnormal phenotypes are provided in Supplementary Table 1.

**Table 1:** The top 20 abnormal phenotypes in term of AUC, together with the tumor type that provides the best predictor

| phenotype | tumor type | AUC |
| --- | --- | --- |
| Round face | STAD | 0.760 |
| Cafe-au-lait spot | BRCA | 0.746 |
| Dental malocclusion | STAD | 0.742 |
| Thick eyebrow | UCEC | 0.737 |
| Abnormality of the hairline | BLCA | 0.735 |
| Short attention span | STAD | 0.735 |
| Attention deficit hyperactivity disorder | STAD | 0.734 |
| Abnormality of the metatarsal bones | STAD | 0.733 |
| Congenital abnormal hair pattern | BLCA | 0.732 |
| Abnormality of the maxilla | STAD | 0.732 |
| Hypermetropia | STAD | 0.731 |
| Misalignment of teeth | STAD | 0.729 |
| Hyperactivity | UCEC | 0.729 |
| Small face | UCEC | 0.729 |
| Abnormal size of the palpebral fissures | STAD | 0.728 |
| Joint hypermobility | UCEC | 0.727 |
| Hypoplasia of the maxilla | STAD | 0.726 |
| Aplasia/Hypoplasia involving the metacarpal bones | STAD | 0.726 |
| Low posterior hairline | BRCA | 0.725 |
| Abnormality of the posterior hairline | BRCA | 0.725 |

### 2.5 Validation on new phenotype-gene associations

To validate the usefulness of the TSM-based model in predicting new gene-phenotypes associations we took advantage of a major update of the HPO annotations that took place between October 2015 (annotations that were used in training the models) and the build downloaded in July 2016. A total of 28765 annotations were added to the database, 25120 of which associating a gene and a phenotype included in our models.

We thus selected, for each phenotype with new gene associations, the model based on the TSM in the cancer type that gave the best prediction in the training data, and recorded the rank of the newly associated genes in the prediction. As a comparison we repeated the procedure after randomly relabeling 100 times the genes. These results show that TSM-based models are useful in predicting gene-phenotype associations that were not included in the training set. Fig. 3 shows the rank distribution of true new associations (green bars) and the same after random relabeling (black dots).

**Figure 3:**
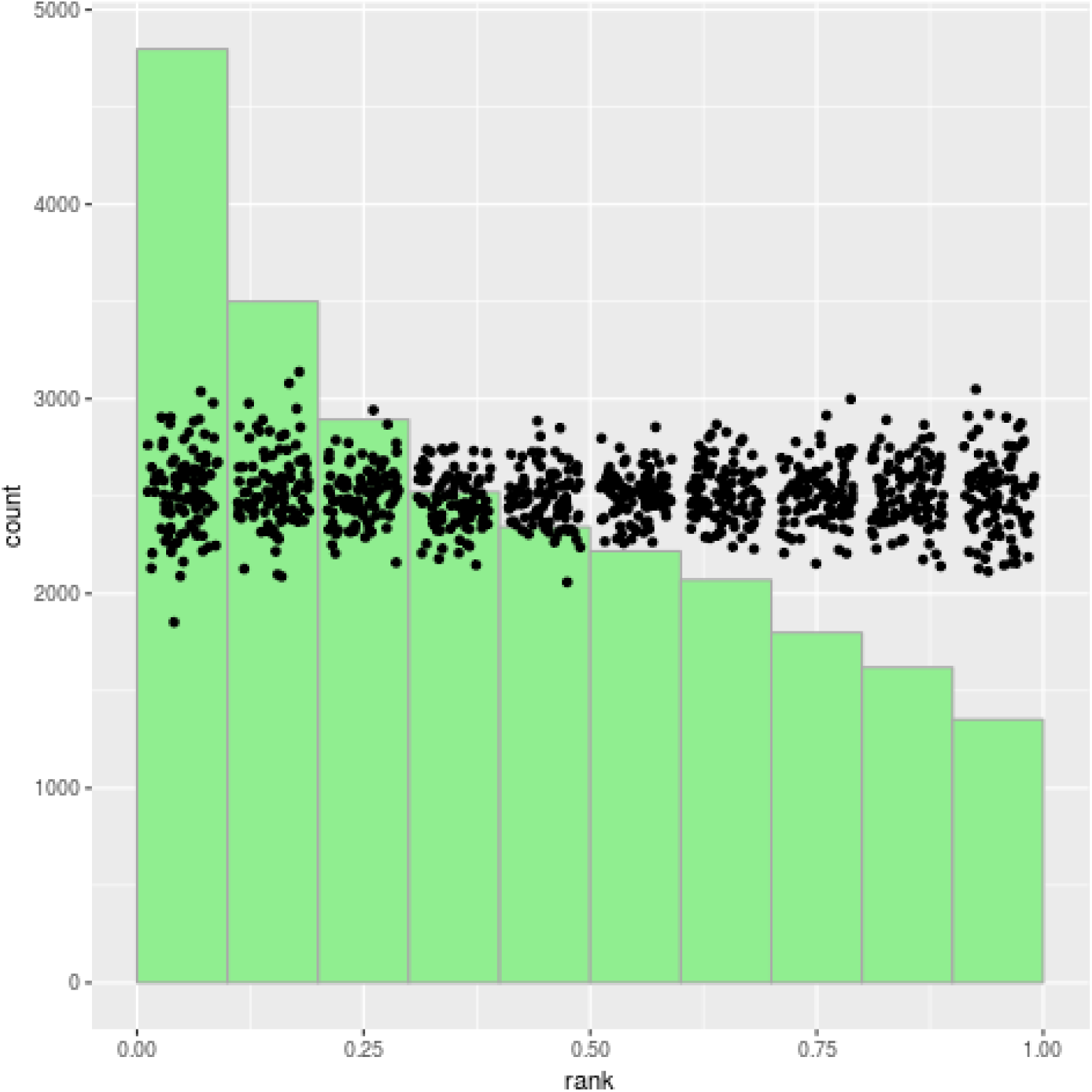
Rank of gene-disease associations not used in training the model. The dots represent the distribution of the same ranks after 100 random relabeling of the gene names

### 2.6 Enrichment analysis of top-ranking genes

We used Gene Set Enrichment Analysis (GSEA) [9] to determine the functional enrichment of the genes predicted by TSM to be involved in abnormal phenotypes, limiting the analysis to Gene Ontology gene sets. When considering the rank of the genes in the TSM predictions, each of the 1007 HPO abnormal phenotypes we study corrsponds to one of 22 non-redundant rankings of the genes (the ranking for each phenotype depending on the best-performing tumor type for that phenotype and the sign of the prediction, namely whether greater or lower TSM corrlates with involvment in the phenotype).

Many gene sets are enriched in more than one ranking: Table 2.6 shows the gene sets falling into the top 20 most enriched in 10 or more rankings (after removing a few redundant gene sets: the complete list of recurrent gene sets is in Supplementary Tables 2 and 3).

**Table 2:** Gene Ontology gene sets showing significant enrichment in several TSM predictors

|  |  |
| --- | --- |
| MEMBRANE DEPOLARIZATION DURING ACTION POTENTIAL | 19/22 |
| HOMOPHILIC CELL ADHESION VIA PLASMA MEMBRANE ADHESION MOLECULES | 19/22 |
| GLUTAMATE RECEPTOR SIGNALING PATHWAY | 18/22 |
| LIPID TRANSLOCATION | 16/22 |
| NEURON RECOGNITION | 11/22 |

These results suggest that the same mutations disrupting intracellular signaling can allow proliferation and invasion by cancer cells on one hand and jeopardize the delicate cell-cell communication needed during development. Specifically, intracellular signaling in the central nervous sytem appears to be most often involved in such disruption.

### 2.7 From phenotypes to diseases

We asked how the cancer TSM would perform in predicting genes associated to genetic diseases, rather than individual phenotypes. The HPO site provides a table associating a list of phenotypes to each ORPHANET disease. A disease gene predictor can thus be obtained by suitably combining the predictions for its associated phenotypes. We chose to aggregate the phenotype predictors by rank products: the score of a gene as a candidate for a disease is the geometric mean of the ranks of the gene as a predictor of the phenotypes associated to the disease. The performance of such predictor on true associations is shown in Fig. 4, where the dots represent the performance after gene randomization as described above.

**Figure 4:**
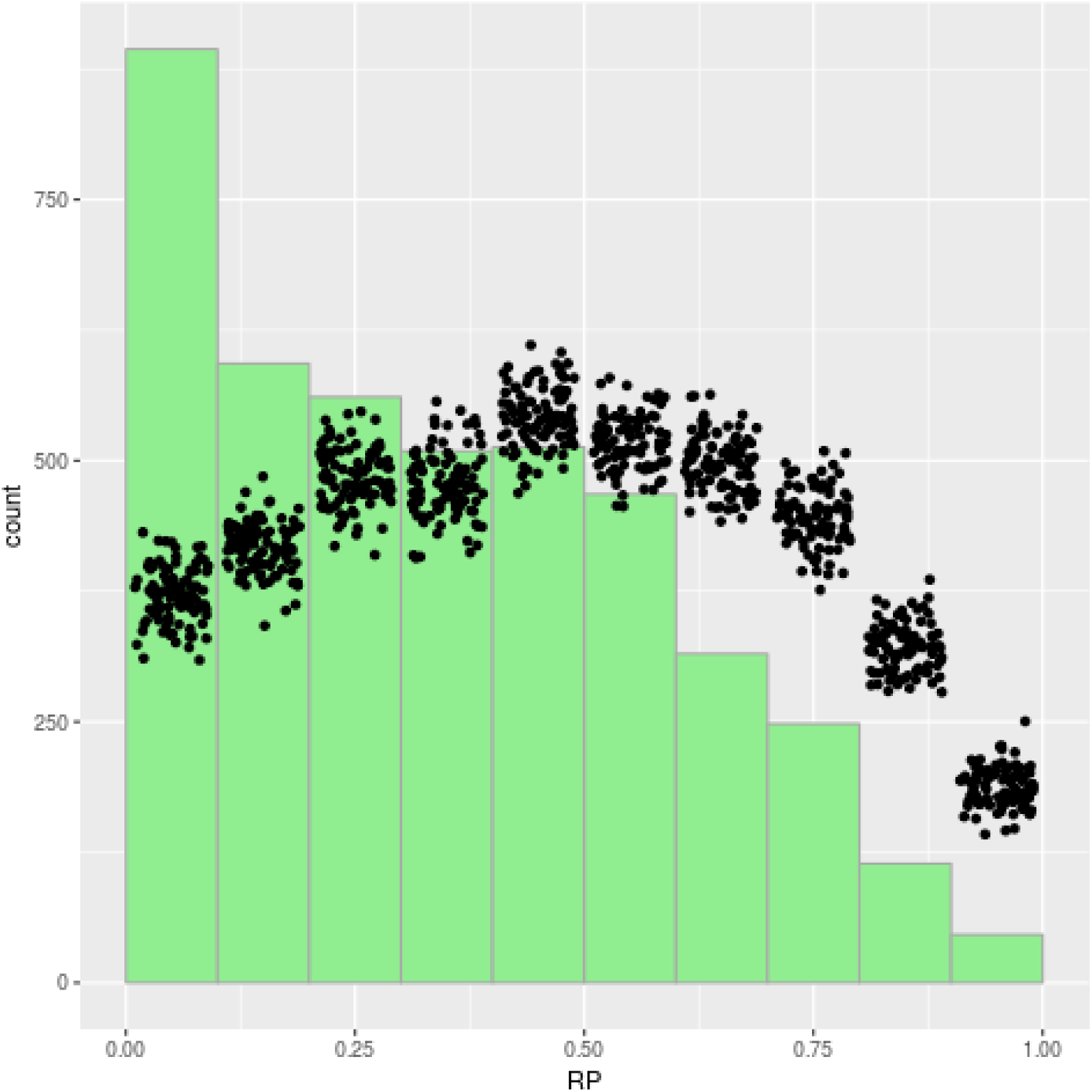
Rank product of true disease genes associated to Orphanet terms

The result is robust when applied only to disease-gene associations established after 2015, and thus not used in training the phenotype models (Fig. 5)

**Figure 5:**
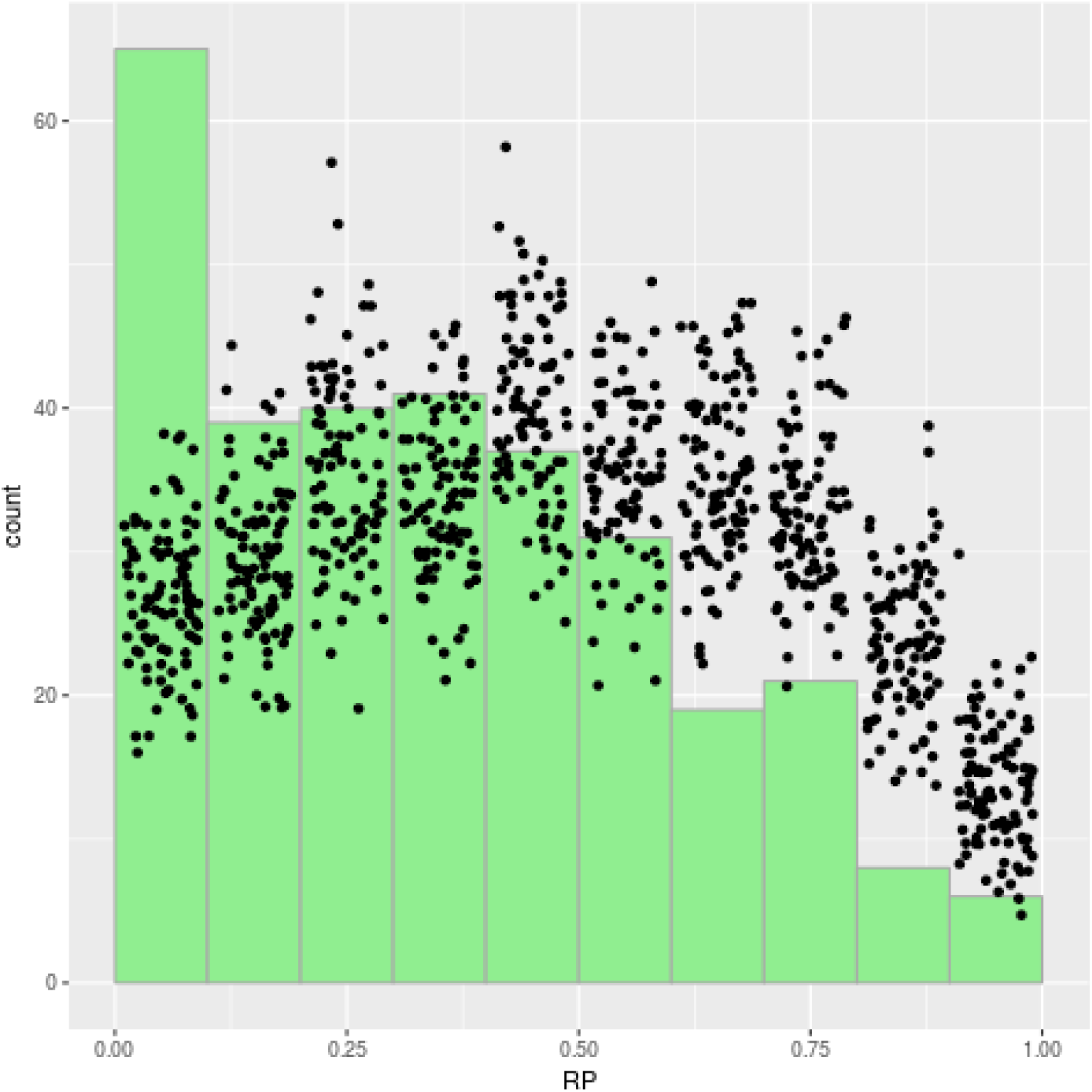
Rank product of true disease genes associated to Orphanet terms, limited to associations established after 2015

## 3 An example: prioritizing *de novo* variants

To show that the TSM-based predictors are useful in practice when prioritizing variants, we considered a recent analysis of *de novo* variatnts in Autism Spectrum Disorder (ASD) [10]. The authors divided such variants in two classes according to whether they are found in the ExAC database of non-diseased exomes (class 2 variants) or not (class 1 variants). Class 1 variants are more likely to be causally involved in the disease.

We classified the genes in which *de novo* variants were found in children affected by ASD into two classes: those with at least one class 1 variant (“class 1 genes”) and those without (“class 2 genes”), and compared their ranks in the best predictor for phenotype “Autistic behavior”. The results are shown in Fig. 6, and show that the TSM-based predictor significantly prioritize Class 1 over Class 2 genes:

**Figure 6:**
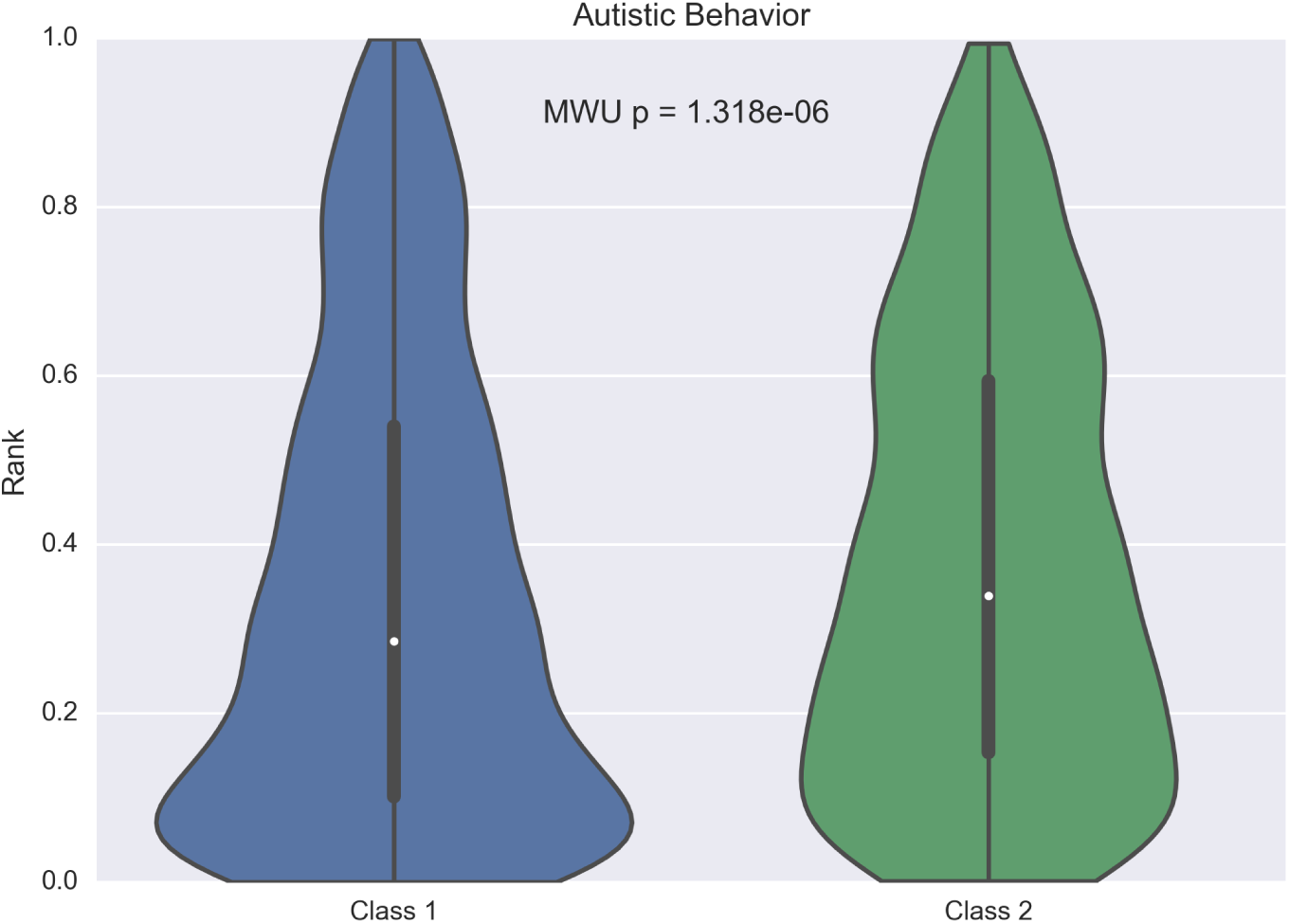
Rank of genes for which *de novo* mutations were found in autistic children. Class 1 genes are those for which not all variants found in affected individuals are found in ExAC, and thus more likely to be causally involved in the disease. The *P*-value is obtained with a Mann-Whitney *U* test

The same analysis can be done on other diseases for which the same data are available in the Supplementary material of the same paper. For intellectual disability and schizophrenia we find the same significant prioritization of class 1 genes, while for congenital heart disease we find the same trend but without statistical significance (Suppl. Fig. 1)

## 4 Discussion and conclusions

We have shown that the large-scale assessment of somatic mutations in tumors is not only useful in understanding the genetics of cancer, but also in an unexpected context, namely in predicting the *germline* mutations responsible for phenotypes and diseases. Genes that are recurrently mutated in tumors are more likely to be involved in abnormal phenotypes when mutated in the germline. This connection had been suggested based on specific cases, but here we have shown that it is generally true in a statistically controlled way.

These results have practical implications and raise conceptual issues. From the practical point of view, they suggest that TSM profiles could be profitably integrated with other sources of information (germline variation patterns, functional annotation, biomolecular networks, etc.) in developing tools for the prioritization of disease genes.

Conceptually, we should ask what is the origin of this correlation between the frequency of somatic mutations and the phenotypic effects of mutations in the germline. We propose two possible mechanisms, not mutually exclusive. First, both somatic mutations in cancer and germline mutations leading to abnormal phenotypes must be compatible with viablility at the cell level. Therefore cancer genomics provides us with a catalog of mutations that are compatible with cell-level viability and thus potentially involved in abnormal phenotypes compatible with life. Indeed, as expected, TSM negatively correlates with essentiality at the cell level as determined in two studies [11, 12] (resp. *P* = 8.5 · 10^−13^, 8.0 · 10^−10^, logistic regression).

However this cannot be the whole story since, as shown in the case of *de novo* mutations involved in autistic behavior, intellectual disability and schizophrenia, TSM is able to predict the most likely causative mutations even among a set of mutations that were actually found in individuals, and hence compatible with cell-level viability. Since TSM represents the recurrence of mutations in cancer we can assume that it measures the growth advantage conferred to the cells carrying them. These could be the same mutations that, in the germline, significantly alter the balance between cell types during development, leading to abnormal phenotypes.

Such relationship with development is confirmed by looking at the predictive power of TSM-based predictors as a function of the age of onset of diseases. In Fig. 7 we show the rank of true disease genes as predicted by rank products for diseases of varying age of onset. We can notice that the performance of the TSM-based prioritizer is indeed maximal for diseases for which the onset is antenatal while the performance decreases for diseases manifesting themselves in childhood and adolescence. Surprisingly, the predictive power becomes higher for adult-onset diseases. Such connection with development is also suggested by a functional enrichment analysis of the genes that are top-ranked in the prediction for phenotypes, where intracellular signaling, in particular in the central nervous system, appears to be often disrupted in both cancer and genetic diseases.

**Figure 7:**
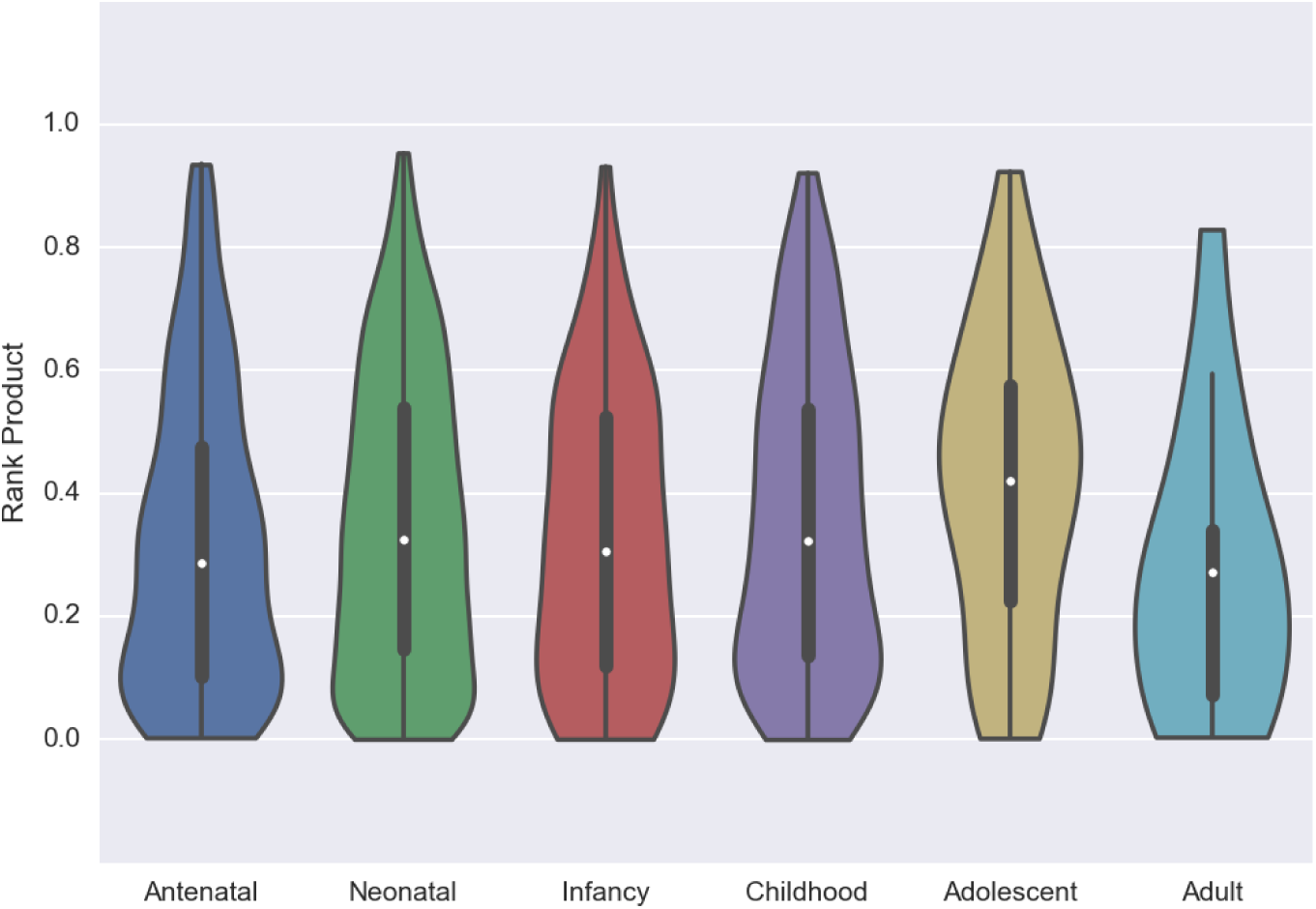
Rank product of disease genes associated to ORPHANET diseases as a function of the age of onset

## 5 Methods

### 5.1 Somatic mutations profiles

Somatic mutation profiles for primary tumor samples of 29 cancer types, summarized at the gene level, were obtained from the UCSC Cancer Browser on November 23, 2015. When several gene-level mutation datasets were available (corresponding e.g. to different analysis pipelines), we based our analysis on the dataset including the largest number of samples, to which we added the samples present only in smaller datasets, considered in decreasing order of sample number. Only protein-coding genes were considered (i.e. those associated with one or more Uniprot ids in the org.Hs.eg.db package of Bioconductor, version 3.4.0). In this way we obtained mutation profiles for 18499 genes in 29 cancer types. Only non-silent mutations were considered in our analysis, according to mutation impact reported by TCGA

### 5.2 Gene-phenotype associations

Gene-phenotype associations were obtained from the Human Phenotype Ontology web site on October 9 2015. We only considered the phenotypes that are direct descendat of the term “abnormal phenotype” (HP:0000118) and excluded the “neoplasm” term (HP:0002664) and all its descendants. Genes associated to a phenotype were associated also to all its ancestors in the HPO graph. Finally we limited the analysis to 1007 phenotypes associated to more than 50 genes.

### 5.3 Logistic regression

We fitted a univariate logistic model for each HP phenotype, where the regressed variable is the association between a gene and the phenotype, and the regressor for gene *g* is one of the following (*n*(*g*,*t*) is the number of samples of cancer type t where *g* is mutated)

- the number of samples in which *g* is mutated, summed over all cancer types (which we refer to as TSM, Total Somatic Mutations) 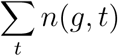
- the log-transformed number of samples in which *g* is mutated, summed over all cancer types 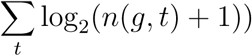
- the mutation frequency of g summed over all cancer types 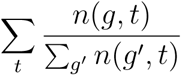
- the log-transformed mutation frequency of g summed over all cancer types 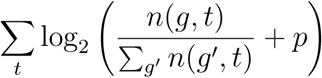
 where *p* is pseudo-frequency equal to 10^−6^. Since all these predictor showed very similar performance on HPO phenotypes we discuss only the first one, which achieves a sligthly higher median AUC.

We also fitted univariate logistic models in which the regressor is the number of mutated samples for a specific cancer type *n*(*g*,*t*).

For comparison we fitted models using predictors based on germline variation frequencies:

- the Residual Variation Intolerance Score (RVIS) determined in [6]
- the probabilities determined in [7] of being loss-of-function intolerant (pLI), intolerant of homozygous LOF (pRec) and tolerant of both homozygous and heterozygous LOF (pNull). We chose pNull for our comparisons since it is by far the best predictor of HPO associations.

Bivariate logistic models were generated to determine if TSM provides independent information on disease genes than what provided by methods based on germline variation frequencies.

### 5.4 AUCs and *P*-values

The performance of logistic models was evaluated using AUCs, and its statistical significance by Mann-Whitney *U* test, in which we compare the values of the model prediction on positive genes (associated to the phenotype) *vs* negative ones. It is worth noting that the for univariate models the AUCs of the logistic model are the same as the AUCs that we would obtain using directly the regressor as predictor, possibly after changing its sign: the only job performed by logistic regression is to determine whether the regressor is positively or negatively correlated with the response.

### 5.5 Predicting disease genes

We obtained the associations between Orphanet diseases and HPO phenotypes from the phenotype_annotation.tab file downloaded from the HPO web site on February 5, 2015. We then built a predictor for each disease by associating to each gene the geometric mean of the ranks of the gene as a predictor of each of the phenotypes associated to the disease (phenotypes with up to 50 genes associated, for which we do not have a prediction, were mapped to their closest ancestor in the HPO graph with more than 50 genes).

Disease gene predictors were evaluated by determining the score distribution of true associations (obtained from the Orphanet web site). As a control we generated randomized predictions through 100 random permutations of the genes for which we have a TSM-based predictions.

### 5.6 Gene Set Enrichment Analysis

GSEA analysis was performed using the Pre-Ranked Analysis module built in the desktop application (v. 2.2.3). Gene lists for each non-redundant prediction were preranked using the prediction rank as natural ordering. Enrichment was evaluated on c5.bp.v5.2 gene list.

## 6 Supplementary Material

Supplementary Table 1: Tumor type with the best predictive power and AUC values for 1007 HPO abnormal phenotypes.

Supplementary Table 2: Gene sets showing recurrent positive enrichment. For each gene set we report the number of non-redundant predictive models in which it is found among the top 20 positively enriched.

Supplementary Table 3: Gene sets showing recurrent negative enrichment. For each gene set we report the number of non-redundant predictive models in which it is found among the top 20 negatively enriched.

**Supplementary Figure 1:**
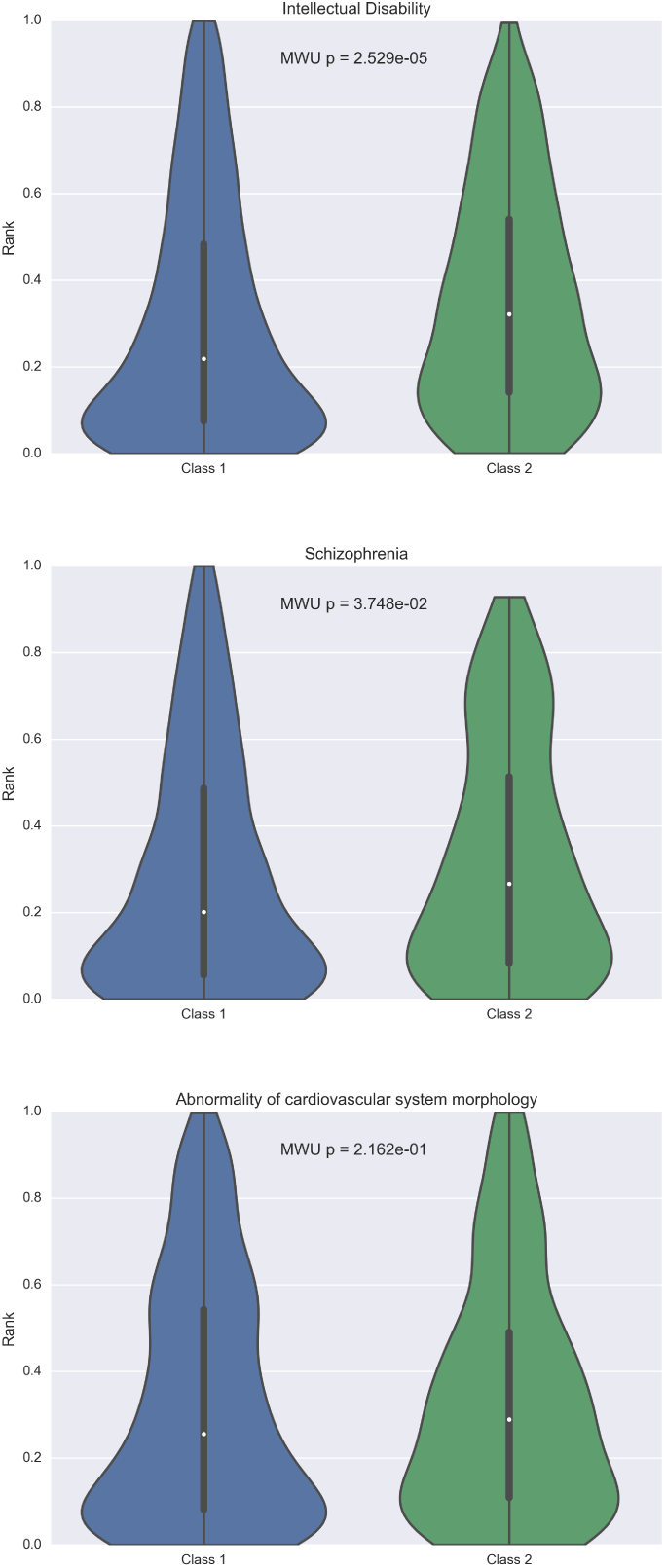
Rank of genes for which *de novo* mutations were found in Intellectual Disability, Schizophrenia and Congenital Heart Disease. Data and classification into classes are from [10]. The following HPO terms were used for the prediction: “Intellectual disability”, “Behavioral abnormality” (the closest ancestor of “Schizophrenia” in the HPO tree for which we have a predictor, since it has more than 50 associated genes) and “Abnormality of cardiovascular system morphology”.

**Supplementary Figure 2:**
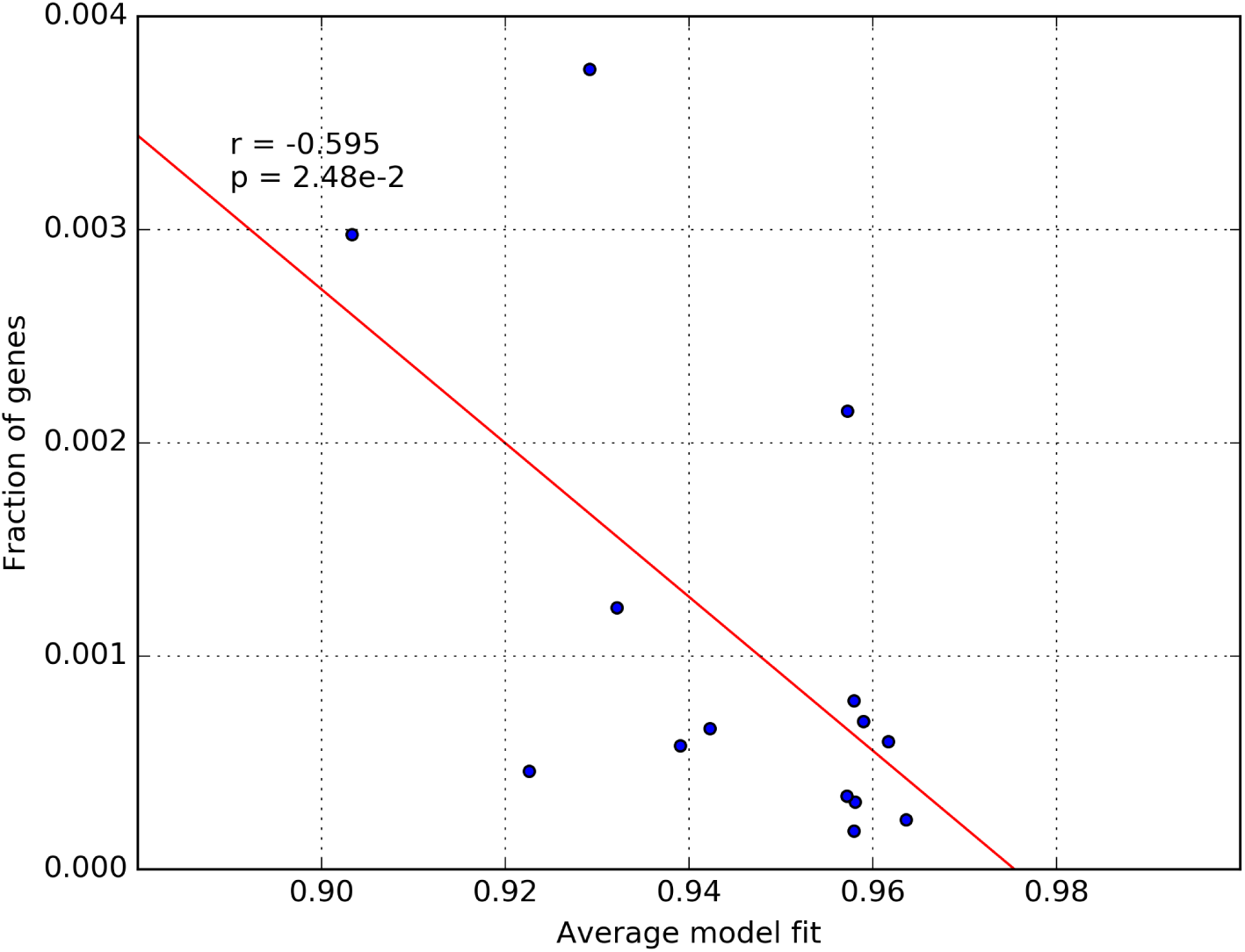
Correlation between model fit to neutral evolution and the fraction of genes having a mutation frequency higher than 1e-3. Values on the *x*-axis are average values displayed in figure 3 from [8].

